# Bacterial Decellularization: Non-Chemical Production of Effective Plant Tissue Bio-Scaffolds

**DOI:** 10.1101/2023.09.27.559696

**Authors:** M.S. Rybchyn, J. Biazik, A.C. Nunez, J. le Coutre

## Abstract

Bio-scaffolds for the cellular agriculture field require to be simple, with low associated costs. Here, a method is described to generate decellularized leaf scaffolds utilizing a simple bacterial process, yielding scaffolds with the capacity to sustain myoblast attachment, growth, and differentiation.

To best develop a minimal cost decellularization process, we aimed to design the key steps of the method to be as “low-tech” as possible and not use chemical or thermal processing to produce the cellular scaffold. Decellularized leaves (DCL) from the black walnut tree (*Juglans nigra*) were successfully produced employing a domestic fish tank with a biological filtration system that supports an active aquatic nitrogen-fixing bacterial population, typically within 3-5 days. Following decellularization, the DCL were devoid of any pulp material as confirmed by scanning electron microscopy (SEM). DCL produced in this way are an effective cellular scaffold, and the C2C12 myoblast cell line was shown to attach, proliferate and differentiate on DCL and maintain viability up to 3 weeks post-seeding. Differentiated cellular material grew extensively over the DCL veins and larger differentiated cellular structures extended between individual DCL veins.

The data presented provide a proof of concept for an inexpensive, simple, and chemical-free method for leaf decellularization, which supports myoblast attachment, growth, and differentiation. The technology provides clear applications for the cellular agriculture field where cost reduction, scalability, and simplification of established laboratory processes, such as bio scaffold production, is a key factor.

## INTRODUCTION

Technologies to produce cell-cultivated meat, i.e., growing edible meat from an animal derived biopsy, are maturing both at industrial and at academic organizations (1,2). With the regulatory approval for cell-cultivated meat products by different companies in different global geographies, and with further applications by additional companies pending approval, the concept of growing meat without the animal has come one step closer to reality (3–5).

Decellularization of plant leaves for the purpose of creating bio-scaffolds for cellular growth is a well described process that normally involves the use of anionic detergents such as sodium dodecyl sulfate, oxidizing chemicals such as bleach and heating of up to 60-70°C to achieve complete decellularization (6–8). Typically, decellularization processes occur under measured laboratory conditions, controlling effectors of the process such as pH, osmolarity and temperature. While these processes have mostly been optimized for tissue engineering applications, bio-scaffolds for the cellular agriculture field should be comparatively simple, with lower associated costs, yet with a high degree of scalability.

Additional requirements are to be met for making a specific scaffolding technology conducive to cellular Agriculture and in particular to produce cell cultivated meat. The bio-scaffold should not harm cells originating from the tissue of choice, for example, muscle cells. Ideally, the bio-scaffold might be able to support growth and differentiation of the target cells in three dimensions. The bio-scaffold should not be animal based in origin and it should be cheap to produce at a scale to competitive with traditional meat production (9).

Moreover, the scaffold must be edible and finally, it will be desirable to have the bio-scaffold provide appealing organoleptic features, most notably texture and possibly taste and nutritional value as well.

Understanding and describing processes at the micro-scale to select favorable initial conditions for meaningful scaling is of critical importance in the cellular agriculture field. It is generally accepted, that the principal components of meat can be produced using a biotechnological approach based on tissue culture. Since the ground-breaking presentation a sdecade ago of a beef patty by Mark Post in 2013 (10), several other milestones have been achieved to show that the technology is feasible in principle. However, it remains unclear if the field will ever achieve an economy of scale, as it would be required to meaningfully provide alternatives to animal-based meet.

Scaling of biological manufacturing processes involves two critical elements. First, it is important to understand the selected process sufficiently well at the micro scale to make sure that any upscaling activity, such as increasing the volume of bioreactors, will not just scale a problem that has not been identified or addressed. Second, unlike increasing the speed or performance of microelectronics, it is becoming evident in the field, that biological processes cannot just be increased in volume or speed, and that novel and unknown phenomena occur at scale. Certainly, there is no such thing as Moore’s law in biology.

Both scaffolding and scaling of cell-based meat materials are connected and intertwined in several ways. Since scaling of cell-based meat to commercial quantities might remain elusive still for some time, many players in the field are using bio scaffolding to obtain larger volumes. Scaffolding of edible myoblast material not only will impart texture and three-dimensionality; it also might provide additional nutrients. It is foreseeable, that in the medium-term future we might be looking at a product spectrum ranging from animal-based meat via all kinds of mixtures between cell-based and plant-based materials, to purely plant-based meat analogues.

Here, we set out to explore the first point, i.e., to understand critical events of myoblast/scaffold interaction at the micro scale. We investigate the utility of decellularized leaves to serve as a scaffolding material for myoblast cells.

The feasibility of ramping up bacterial decellularization of leaves at large scale will need to be tested and established further. However, as this process is based upon live bacteria as the active biological system – and not eukaryotic cells – it is likely that enough material can be produced in order to feed into a cellular agricultural process.

## METHODS

All consumables were purchased from Merck KGaA, Darmstadt, Germany and/or its affiliates unless otherwise stated in the text below.

### Cell culture

The C2C12 myoblast mouse cell line was obtained from the European Collection of Authenticated Cell Cultures (ECACC). Routine passaging of the cell line was carried out in DMEM(High) supplemented with 10% v/v fetal calf serum (FCS). Use of this media was defined as the “standard conditions”. Cells were maintained in a humidified incubator at 37°C/5% CO_2_. For cell expansion and maintenance of the line, cells were passaged and maintained at < 80% confluence to restrict myoblast fusion. No antibiotics were routinely used for cell culture. To promote the differentiated myotube phenotype of the C2C12 line, once seeded on to the DCL structure media was changed to DMEM (high) containing 2% v/v horse serum (HS, ThermoFisher Scientific, MA, USA). Use of this media was defined as “differentiation conditions”.

### Decellularization of walnut leaves

Walnut leaves (Figure 1A) were taken directly from a ∼10-year-old *Juglans nigra* tree. Washed leaves were placed in a domestic fish tank, maintained at 25-27°C, which contained circulating *Nitrosomonas* and *Nitrobacter* bacteria that were continually supplied through an EHEIM classic 250 external filtration system (EHEIM, Deizisau, Germany). While these bacteria are functionally present to reduce ammonia levels in the water, they also effectively break down biological material such as leaf pulp. The tank also contained freshwater shrimp (*Neocaridina davidi*) and Malaysian trumpet snails (*Melanoides tuberculata*) which also likely contributed to leaf pulp removal. Walnut leaves typically decomposed within 3-5 days (Figure 1B). Following decellularization, leaves were either submerged in 80% v/v ethanol (overnight/RT) or autoclaved under steam (121°C/15 min) to sterilize.

**Figure 1.**
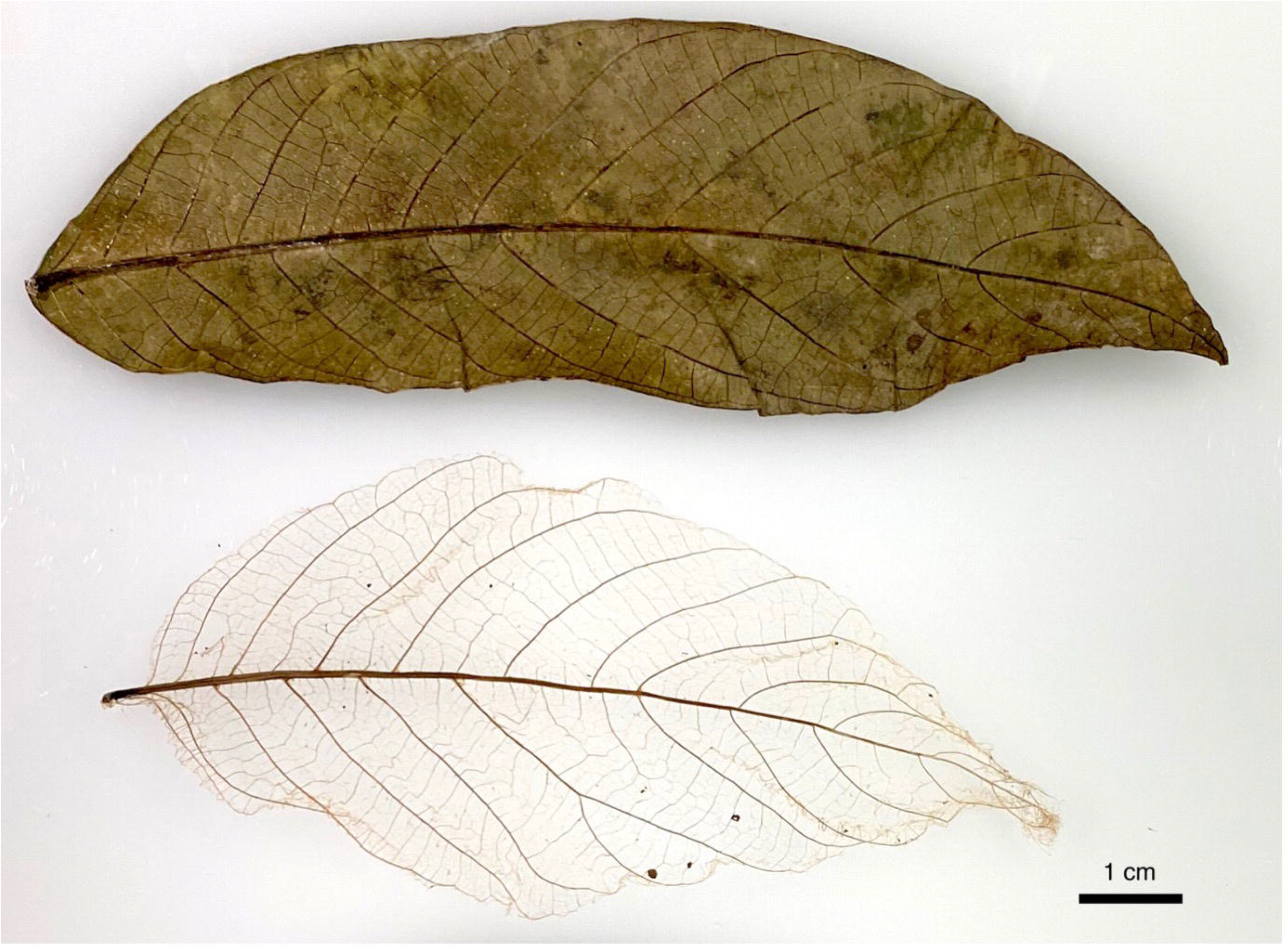
Bacteria effectively decellularize walnut leaves in an aquatic environment. Examples of walnut leaves (*Juglans nigra*) pre-(top) and post-decellularization (bottom) are shown as photographs. Walnut leaves were placed into a fish tank comprising a “living filter” system that dispersed nitrogen-fixing bacteria throughout the tank. Bacterial decellularization took place over several days until complete (typically 3-5 days), at which time leaves were removed and air dried. No additional chemicals, heating or processing were required to achieve complete decellularization. Photography by J. le Coutre.

### Cell seeding onto walnut leaves

C2C12 myoblasts (1x10^6^) or control media were seeded directly onto sterilized DCL of roughly equal size (∼1.5-2 cm^2^) in 200mL of growth media in 6-well plates. A small volume of media was used at a high cell density to promote cell attachment to the structure. After incubation for 1 h (37°C/5% CO_2_) the media volume was increased to 2 mL and incubated overnight (37°C/5% CO_2_). The seeding protocol was repeated twice again on days 2 and 3 post-seeding. At 7 days post-seeding the plate was placed under differentiation conditions and incubated for 2 more weeks. Media changes were performed every 2 days or every day toward the end of the 3-week period due to more rapid media acidification. Penicillin and Streptomycin were used at recommended concentrations throughout this experiment where DCL was used.

### Phase contrast images

Phase contrast images of DCL with or without cells in 6-well plates under differentiation conditions were acquired using the Zen lite software application on a Zeiss Primovert inverted light microscope, with an attached Zeiss Axiocam 105 color camera (Carl Zeiss AG, Oberkochen, Germany).

### Cell Titer Blue assay for cell viability

Following removal of DCL to a new 6-well plate to prevent signal from cells attached to well plastic, the viability dye Cell Titer Blue (CTB, Promega, WI, USA) was added at 10% v/v to wells containing DCL with or without seeded cells, or to media only as a control for 2 h (37°C / 5% CO_2_). Following transfer to black well plates, the fluorescent signal (560ex/590em) was measured on a CLARIOstar plate reader (BMG Labtech, Ortenberg, Germany).

### C2C12 detergent based lysis for bicinchoninic acid assay and western blot

Following removal of DCL to a new well plate to prevent signal from cells attached to well plastic, RIPA lysis buffer (25 mM Tris-HCl pH 7.6, 150 mM NaCl, 5 mM EDTA, 1% Triton X-100, 1% sodium deoxycholate, 0.1% SDS) was added and the solutions were sonicated using the microtip of a Branson 250 Analog Sonifier (output control set to “1”, 50% duty cycle, 20 pulses) to facilitate the release of detergent-soluble cellular protein. The cell lysis solution was centrifuged (10,000 rpm/10 min/RT). A fixed volume of soluble cellular lysate or known concentration of bovine serum albumin (BSA) was used to determine the detergent-soluble protein concentration by Bicinchoninic acid assay. Absorbances of reacted samples were quantified using a CLARIOstar plate reader (562 nm).

### Western blot analysi

Detergent soluble protein (∼10mg) was subjected to SDS-PAGE using a 4-20% Mini-PROTEAN TGX acrylamide gel on a Mini-PROTEAN Electrophoresis cell (Bio-Rad Laboratories, CA, USA). Electrophoretically separated proteins were transferred to Immobilon-P PVDF membrane (25V/RT/overnight). Membranes were blocked using 5% w/v BSA in 10 mM Tris (pH 7.6), 150 mM NaCl containing 0.05% v/v TWEEN20 detergent (TBS-T) for 1 h at RT. Membranes were incubated overnight with primary monoclonal anti-Myosin IIa (clone E7Y9O) at the recommended concentration in blocker (Cell Signaling Technology, MA, USA) followed by anti-rabbit IgG HRP linked secondary antibody at 1:10,000 in TBS-T for 1h at RT (antibodies were from Cell Signaling Technology, MA, USA). Protein bands were detected by exposing the membrane to SignalFire™ ECL substrate (Cell Signaling Technology) on a ChemiDoc Imaging System (Bio-Rad).

### Scanning Electron Microscopy

Following a brief wash with an excess of PBS (pH 7.4), DCL with or without seeded cells were fixed (overnight/4°C) in 2.5% w/v glutaraldehyde, 0.2 M sodium phosphate (pH 7.4). Following rinsing in 0.1 M sodium phosphate (pH 7.4), samples were dehydrated in a graded series of ethanol (30-100% v/v). Samples were then dehydrated using increasing concentrations of hexamethyldisilizane (HDMS) and left to air dry in a final 100% solution of HDMS. Several regions from replicate DCL were chosen at random and mounted on to SEM stubs, platinum coated and viewed using an FEI Nova NanoSEM (Oregon, USA) operating at 5 kV at which time images were acquired.

For presentation purposes and to aid visualization we converted the native SEM greyscale images to an RGB format. Greyscale SEM images in tiff format were imported to Adobe Photoshop (Adobe Inc., CA, US) and converted to RGB format. The neural filter “Colorize” was used to process the greyscale image to a coloured reproduction with the following manually assigned colours in hexadecimal format – “dd2424” (visually red) for cellular deposits; “eea11b” (visually brown) for DCL; “000000” (visually black) for the slide background. Cellular material was clearly identifiable compared to DCL due to their visually distinct structure, the presence of cellular characteristics such as microvilli on cellular deposits. These differences are clear by visual comparison of the negative control in the absence of cells (Figure 2) to cell-seeded DCL (Figures4-7).

**Figure 2.**
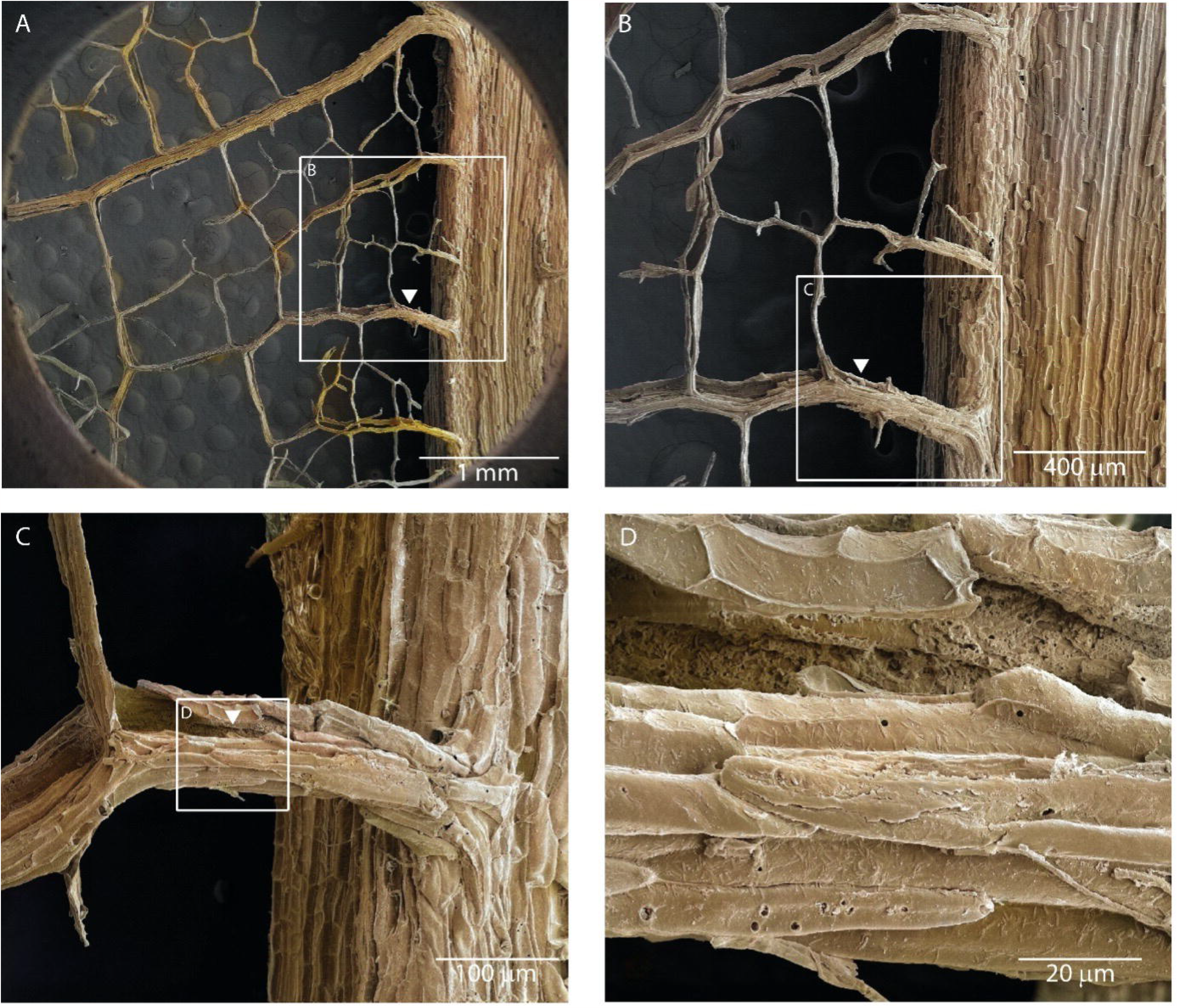
Scanning electron microscopy (SEM) of biologically decellularized walnut leaves (DCL) reveals structural features amenable to promote myoblast attachment. (A-D) SEM at increasing magnifications shows that the DCL stem and veins have an uneven surface and are spatially arranged so that individual veins are typically anywhere from 0.1 to 1 mm apart. In some instances, hollow or “pocket” structures were discernable in the DCL, presumably where the xylem/phloem vascular bundle originally existed (white arrowhead). Insets are shown (A-C) to show the relative location of increasing magnification. SEM magnifications were as follows (A-73x; B-181x; C-648x; D-3230x). Colouring of the original greyscale images is described in Methods. The original greyscale images with associated metadata from the SEM are shown in Figure 1 Supp.

Throughout, a conservative approach has been pursued to assign any region of the image as being cellular material for the purposes of assigning colour, and if there was any ambiguity or both cellular (red) and DCL (brown) was present in an image region then the region was coloured as DCL (brown). For this reason, a minute amount of cellular material in the presented images, most often at the boundary between DCL and cellular material, may be incorrectly colored. For all figures presented, the original greyscale images are shown in the same figure layout as supplementary data for reader comparison and to provide transparency of the image coloring process.

### Statistical Analyses

GraphPad Prism 9.2 was used for all statistical analyses.

## RESULTS

### Leaf decellularization is achieved in an aqueous, microbial environment without heating or addition of chemicals

Decellularization of a leaf (Figure 1A) taken freshly from a black walnut tree (*Juglans nigra*) was performed via incubation of the leaf in a domestic fish tank for several days, which was utilized as an aqueous environment rich in nitrogen-fixing bacteria. This process was effective as the outer green layer and pulp of the leaves was no longer present typically within 3-5 days, and the veins, midrib and petiole remained intact (Figure 1B). A more detailed examination by SEM at several magnifications shows that the decellularized leaf (DCL) structure was completely devoid of leaf pulp material (Figure 2). At higher magnifications, SEM shows that the outer structure of the DCL veins was generally uneven and had “pocket” structures that we predicted would be effective at trapping individual myoblast cells to permit cellular attachment and growth over the DCL structure (Figure 2, white arrowheads).

### Decellularized leaves are an effective scaffold for maintaining viable mammalian myoblasts

C2C12 myoblasts were seeded directly onto sterilized DCL structures in 6 well plates and maintained for 2 weeks under differentiation conditions. Phase contrast microscopy at this time reveals that mammalian cellular material was localized to and around the leaf skeleton, and extended between the leaf veins (Figure 3A, white arrowhead). This cellular material was not observed by phase contrast on negative controls (Figure 3B).

**Figure 3.**
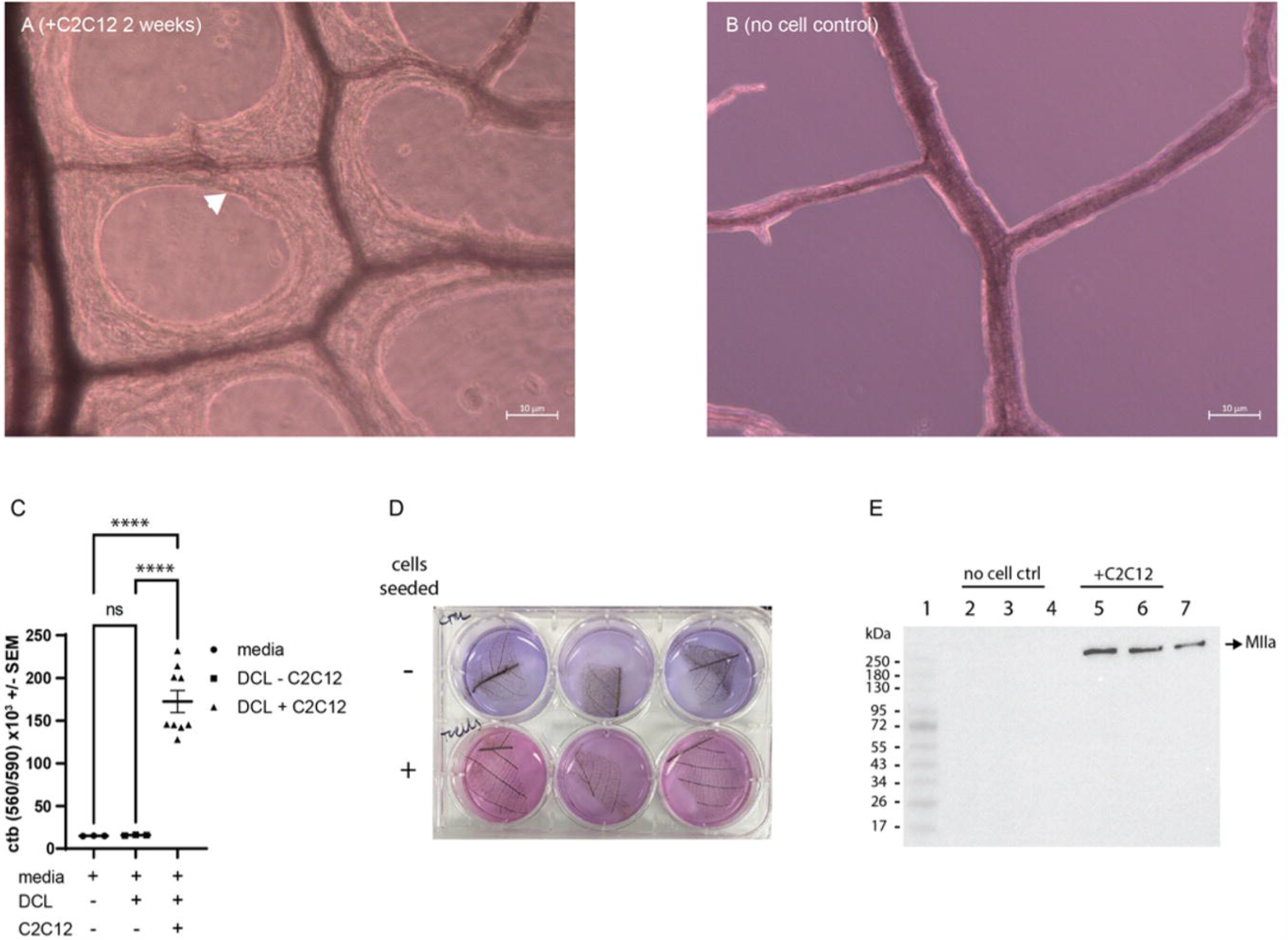
Viable C2C12 myoblasts attached and grew on decellularized leaf structures (DCL). (A-B) Phase contrast images of DCL that (A) had been seeded with C2C12 myoblasts or (B) the no cell-seeded control were acquired at 2 weeks post seeding (scale bars = 10µm). Cellular growth was clearly observed around the DCL veins where cells had been seeded onto DCL (A, white arrowhead). (C) At 3 weeks post-seeding, DCL that had been seeded with C2C12 myoblasts (“C2C12”, n=9), or no cell DCL controls (“DCL”, n=3) were removed from their existing 6 well plates, washed with PBS and placed in new 6 well plates with fresh media, alongside an equal volume of media to serve as a *media only* control lacking DCL (“media”, n=3). The resazurin dye Cell Titer Blue (CTB) was added to detect viable cells. CTB signal was quantified by fluorescence intensity after transfer of equal volumes of the reactant to a black-well plate (λ560_ex_/λ590_em_). “ns” not significantly different; **** p<0.0001 (One-way ANOVA, Dunnett post-test). (D) A photograph showing the visual difference in CTB reaction between DCL that had been seeded with C2C12 myoblasts (+) or without (-), as was quantified in “C”; Photography by M. S. Rybchyn. (E) A western blot showing the expression of myosin-IIa (MIIa) from a detergent-based lysis of DCL that had been seeded with C2C12 cells for 3 weeks prior to lysis (+C2C12) or that present from a no-cell control (no cell CTRL).

Following removal of DCL to new plates to avoid signal from cells attached to the plastic wells, the addition of the resazurin dye Cell Titer Blue established the presence of viable cells attached to the decellularized leaf structures (Figure 3C&3D). No significant difference in fluorescent signal is observed between media and DCL only control without seeded cells (Figure 3C). DCL with seeded C2C12 myoblasts yielded fluorescent readings ∼10-fold higher than control (p<0.0001). Western blot analysis of DCL that was seeded with or without cells reveals the presence of myosin IIa protein, an expected marker of a differentiated muscle cell population that was not detected from the negative control (Figure 3E). An estimation of total cellular protein attached to the leaf structure by BCA assay yielded a value of 2.7 (± 0.5) mg per cm^2^ DCL, and the inability to easily solubilize the differentiated cellular material bound to the DCL structure likely resulted in an underestimation of this quantification.

### Differentiated cellular material grows tightly over and between veins of DCL

SEM analysis at low magnification (∼70x) of cell seeded DCL shows that a high proportion of the DCL was covered with differentiated cellular material at 3 weeks post-seeding (Figure 4). Higher magnification SEM revealed that the cellular material grew extensively both covering the DCL vein structure (Figure 5) and between individual veins of the DCL vein structure (Figure 6). In many of the high magnification SEM graphs presented, individual cells or groups of cells are observed at the boundary of the differentiated cellular material, and on top of the differentiated cell layer in some instances suggesting that cellular proliferation was likely still occurring on the DCL structure up to 3 weeks post seeding (clear example of undifferentiated myoblasts is shown in Figure 5B, white arrow).

**Figure 4.**
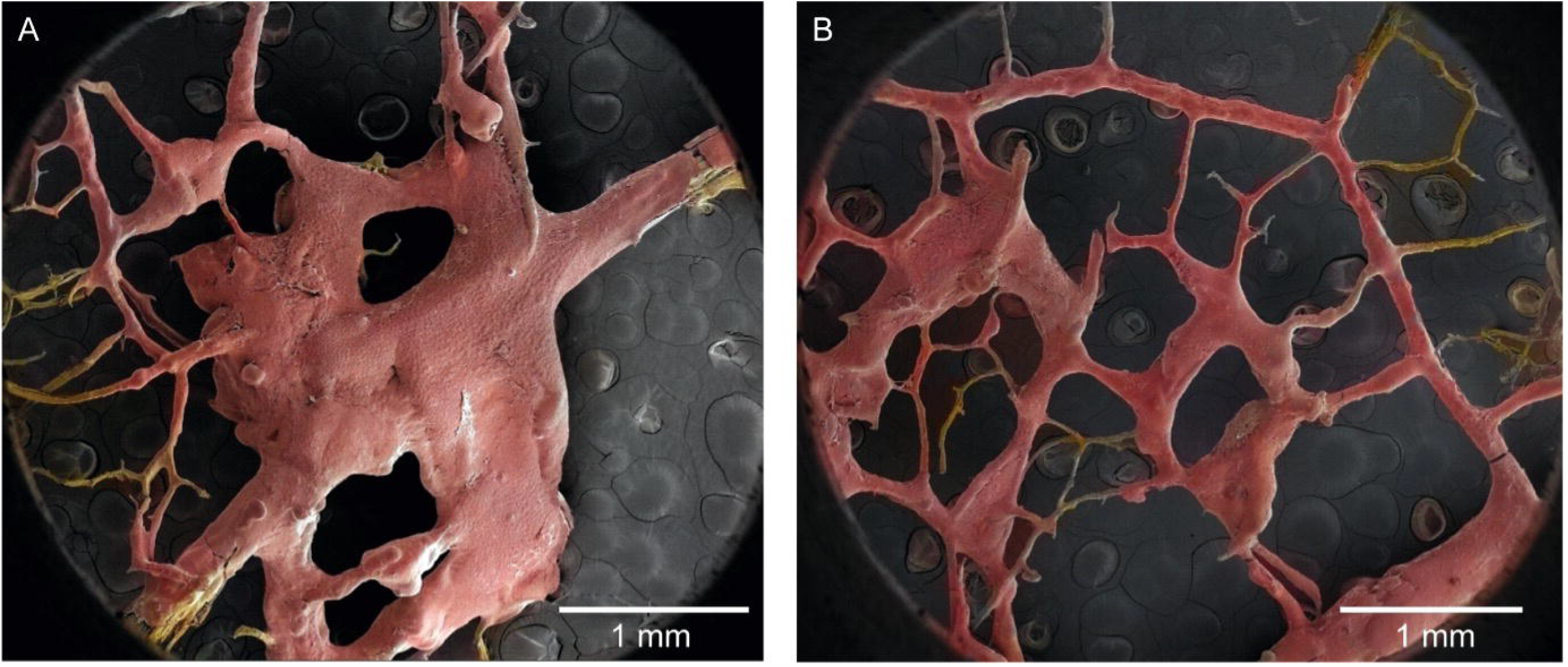
C2C12 grow and differentiate extensively over a large proportion of the decellularized leaf (DCL) structure as revealed by low magnification SEM. (A-B) Low magnification (A-73x; B-71x) SEM images of DCL that was seeded with C2C12 myoblasts and maintained for 3 weeks to promote cellular differentiation. Areas of the image determined to be cellular material are shown as red, DCL without cellular material attached are shown as brown, and the slide background is shown as black. Colouring of the original greyscale images and identification of cellular material is described in Methods. The original greyscale images used to produce the coloured images shown (A-B) with their associated metadata from the SEM are shown in Figure 2 Supp.

**Figure 5.**
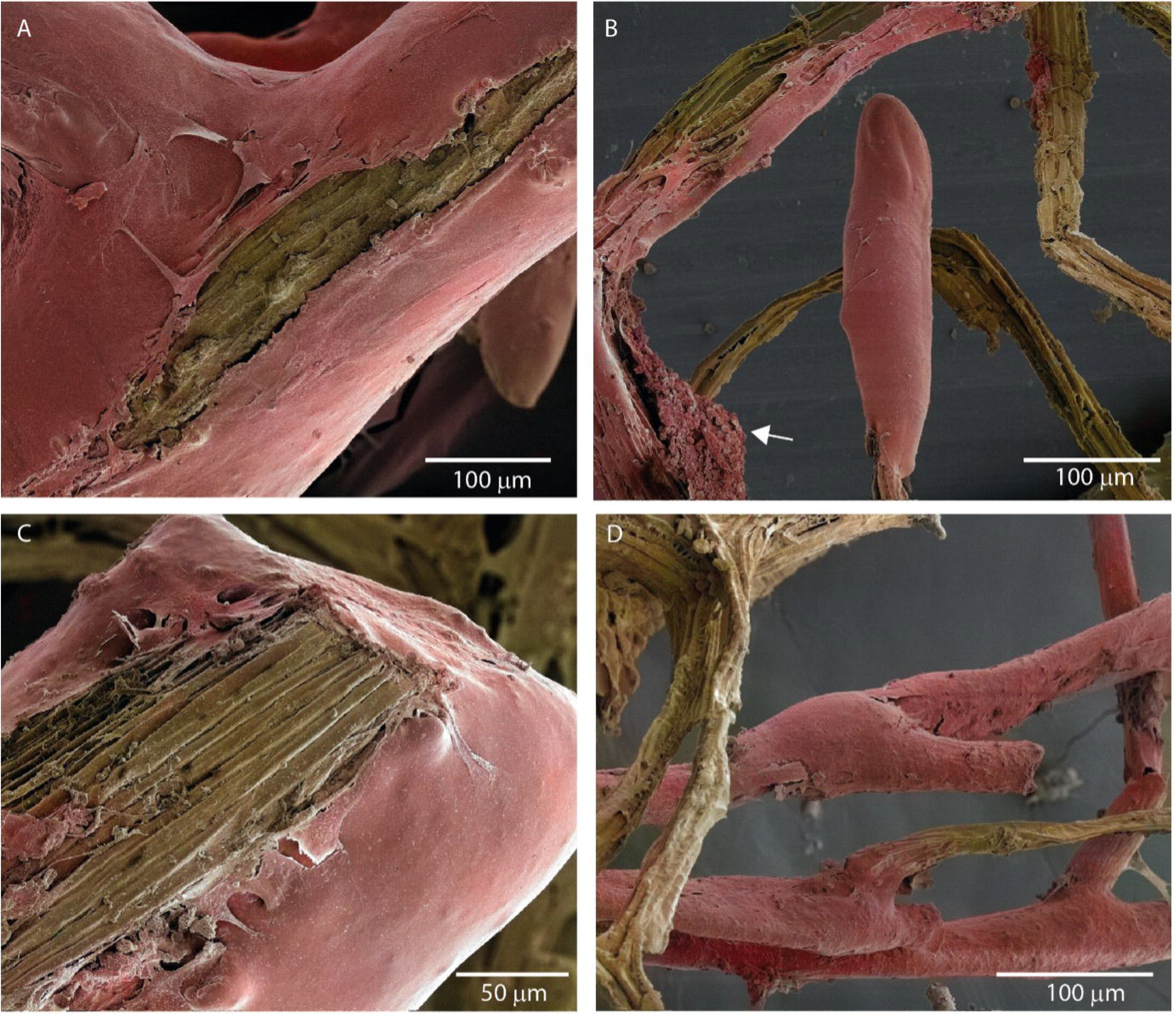
Differentiated cellular material grows “tightly” over the veins of decellularized leaf (DCL) structure. (A-D) SEM images of DCL that was seeded with C2C12 myoblasts and maintained for 3 weeks to promote cellular differentiation. Differentiated cellular material grew tightly over the veins of the DCL. SEM revealed instances where cellular material grew to completely (B and D), or partially (A and C) cover the DCL vein structure (C, white arrow). The presence of undifferentiated myoblasts attached to DCL are indicated. Areas of the image determined to be cellular material are shown as red, DCL without cellular material attached are shown as brown, and the slide background is shown as black. Colouring of the original greyscale images and identification of cellular material is described in Methods. The original greyscale images used to produce the coloured images shown (A-D) with their associated metadata from the SEM are shown in Figure 3 Supp.

**Figure 6.**
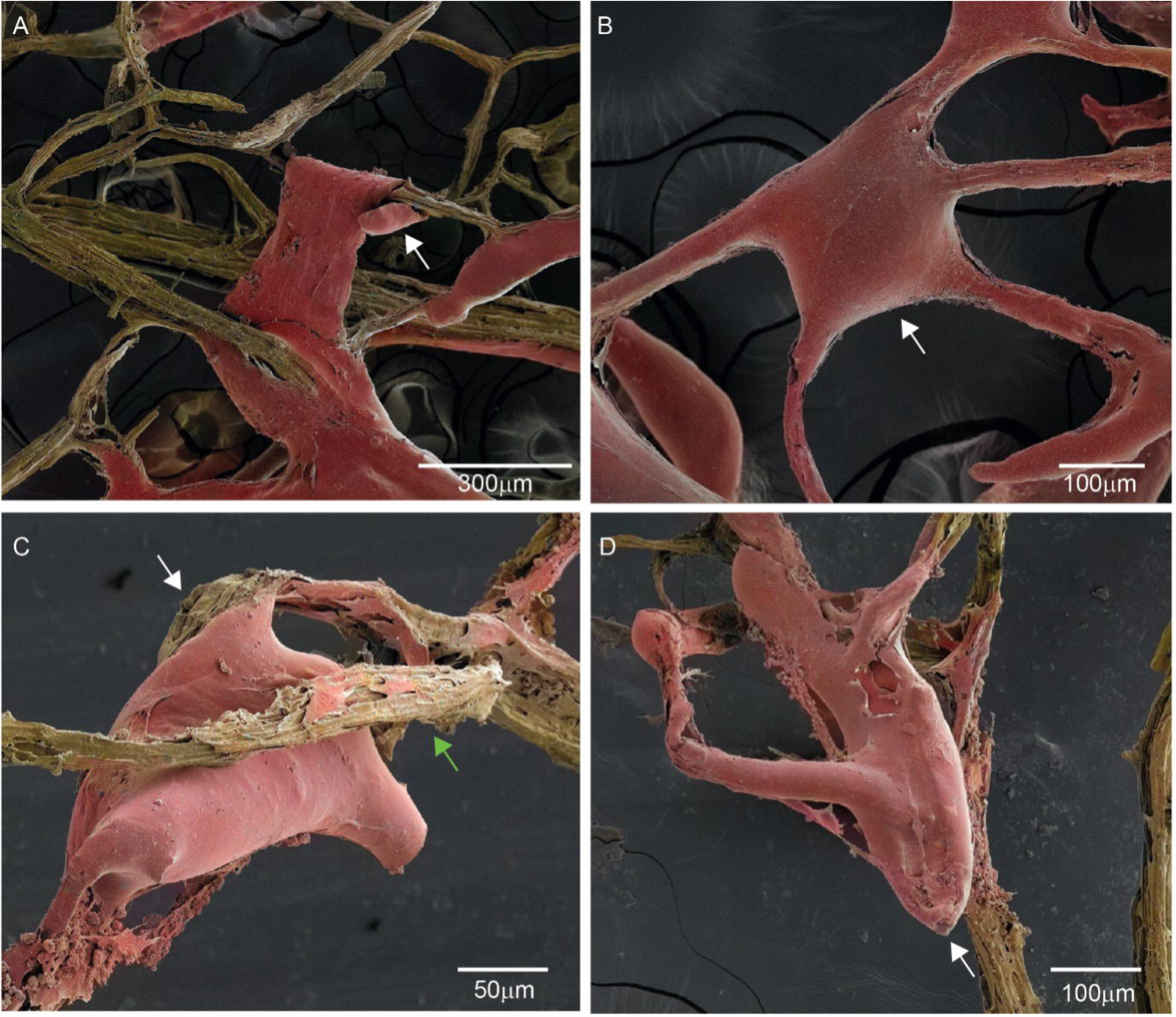
Differentiated cellular material grows between the individual veins of decellularized leaf (DCL) structure. (A-D) SEM images of DCL that was seeded with C2C12 myoblasts and maintained for 3 weeks to promote cellular differentiation. Differentiated cellular material grew between the veins of the DCL producing *de novo* cellular structures. SEM revealed instances where; in (A) a novel differentiated cellular structure grew to be supported at either end by the DCL, as shown by the “looping” of the cellular material over the DCL (A, white arrow); in (B) a *de novo* differentiated cell structure was detected growing between the DCL veins with a central mass (B, white arrow) and “arms” extending to the DCL veins to support the structure; in (C) where the mass of a cellular growth attached to one DCL vein (C, white arrow), was sterically supported by a second vein (C, green arrow); in (D) where a cellular growth was supported by several attachment points to DCL veins and grew in to a large supported mass (D, white arrow). Areas of the image determined to be cellular material are shown as red, DCL without cellular material attached are shown as brown, and the slide background is shown as black. Colouring of the original greyscale images and identification of cellular material is described in Methods. The original greyscale images used to produce the coloured images shown (A-D) with their associated metadata from the SEM are shown in Figure 4 Supp.

### De novo myofibrillar structures are growing from anchor points on the DCL scaffold

SEM reveals *de novo* cylindrical myofibril structures are growing within the DCL structure (Figure 7). These myofibrillar structures were actively growing “outwards” from anchor points on the DCL scaffold (Figure 7A, green arrow). Higher magnification SEM reveals that active cellular remodeling and/or growth appeared to be occurring at the end of certain fibrils suggesting fibril formation was still occurring at 3 weeks post seeding (Figure 7B and 7D). In some instances, fibril structures are found to be fractured (Figure 7C). Fracturing may have occurred during sample processing for SEM, or as a natural process due to other sources of stress to the DCL (*e*.*g*., shear stress). For some of these fractured fibrils SEM indicates a hollow interior structure (Figure 7C), which differed to fibrils that exhibited signs of active growing/remodeling at their ends which comprised a solid interior (Figures 7B and 7D).

**Figure 7.**
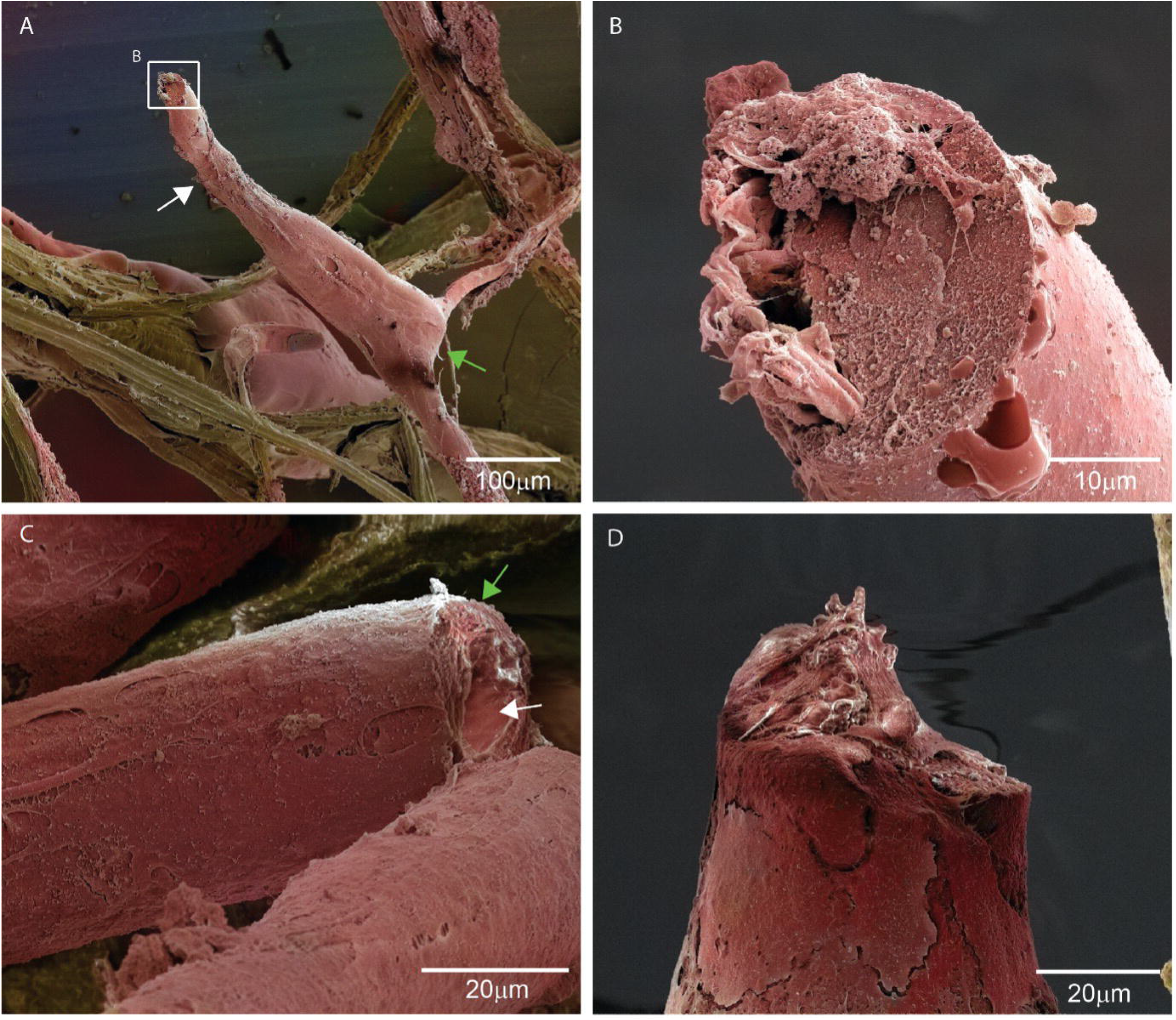
Scanning electron microscopy (SEM) reveals *de novo* differentiated fibril structures are growing from anchor points on the DCL. (A-D) SEM images of DCL that was seeded with C2C12 myoblasts and maintained for 3 weeks to promote cellular differentiation. SEM reveals novel myofibril structures that extended from anchor points on the DCL scaffold. (A) Novel cylindrical myofibril structures (white arrow) extended from anchor points on the DCL (green arrow). (B) A high SEM magnification inset of (A) and (D, a different fibril structure) show active cellular remodeling at the end of the fibril, revealing that myofibrils were still actively forming at the point of fixation (3 weeks post-seeding). The interior of the ends of these actively forming fibrils was solid. (C) SEM of a fractured, pre-formed fibril structure reveals a hollow interior structure (white arrow) and thick cellular outer layers (green arrow), which differed from the active point of fibril formation shown in B and D. Areas of the image determined to be cellular material are shown as red, DCL without cellular material attached are shown as brown, and the slide background is shown as black. Colouring of the original greyscale images and identification of cellular material is described in Methods. The original greyscale images used to produce the coloured images shown (A-D) with their associated metadata from the SEM are shown in Figure 5 Supp.

## Discussion

A key challenge to facilitate the entry of cell-based meat into a consumer accepted, commercially viable, food product is the development of simple bio-scaffolds that do not require a high degree of industrial or lab-based processing. High degrees of processing not only add to cost and waste, but ultimately these processes will be seen as a negative attribute for the consumer. Therefore, natural, readily available scaffolds have an inherent advantage over plastics or other synthetic-based scaffolds. For a comprehensive review on non-animal based cellular scaffolds we direct the reader to the review by Campuzano *et al*.(11).

In this study we describe a biological, rather than the more commonly used chemical methods (12–17) for leaf decellularization that yields an effective bio-scaffold for cell attachment, proliferation, and differentiation. Commonly used chemicals for plant decellularization include sodium dodecyl sulfate (SDS), an anionic detergent, the non-ionic detergent Triton X-100 and bleach, an oxidizing chemical. While these chemicals are all technically biodegradable, their use adds to the cost of the decellularization process, and they are considered harmful to the environment when present at high levels (18–20). As a technical limitation it has been noted that the use of high concentrations of SDS during chemical decellularization can result in higher levels of seeded mammalian cell death (21). This is likely due to SDS molecules remaining bound to the scaffold even with adequate aqueous washing, a common feature of surfactants, and then disrupting mammalian cell membranes following the cell seeding process ultimately leading to an increase in quantified cell death. A biological decellularization process avoids such problems. Our biological decellularization process utilized the nitrogen-fixing bacteria present in fish tanks, which actively remove the pulp material from leaves over approximately 3-5 days. The successful application of a domestic fish-tank for leaf decellularization demonstrates the simplicity and effectiveness of the process, which does not require strict control (other than temperature) or a dedicated technology environment. In terms of scale, a microbial based process for leaf decellularization would likely be more cost-effective and simpler to implement than chemical based processes, that would be more expensive and require more technical equipment and competence by comparison.

We demonstrated that biological decellularization of a leaf yields “pocket” structures in the leaf veins, which were naturally occupied by the xylem/phloem system of the leaf. These access points were large enough for a myoblast in suspension to enter (Figure 2) and were the likely seeding point for myoblast attachment. From here we propose that cellular material is able to attach, proliferate and grow over the entire leaf structure (Figures 4-7). Interestingly, such a “pocket” structure would provide protection from shear stresses encountered in a bioreactor system. Shear stress is a problem for scaffolds in bioreactor systems (refs) and protection from this stress would clearly be advantageous. It is unclear from the current study whether cellular material also expanded throughout the interior hollow system of the leaf. An analysis using resin embedded samples and ultramicrotomy followed by transmission electron microscopy would be necessary to see a cross section of the interior, which was outside the scope of this study. If indeed cellular material also grew on the interior of the structure, then the current analysis by SEM would show an underestimation of the cell/scaffold ratio.

The efficacy of our DCL scaffolds was clear as at 3 weeks post cell-seeding. Under static growth conditions, SEM shows that differentiated cellular material extensively covered our DCL scaffolds (Figure 4). At this stage media acidification was occurring rapidly, with media replacement required every 24 hours, and this was likely a limiting factor for further cell growth in the static system. We propose that in a bioreactor or non-static setup, cellular growth would have continued over and between the veins of the DCL scaffold to an even greater extent. Given that we showed that cellular growth extends between the veins of our DCL scaffolds (Figures 4 & 6) it is likely that cell growth would also extend between individual leaves if they were “packed” into a bioreactor chamber. This, however, remains to be tested.

Simple, cost-effective processes to produce cellular scaffolds that yield high cell/scaffold ratios potentially have applications outside the field of cellular agriculture. Commonly used adherent cell lines for the production of recombinant proteins and vaccines, such as the mammalian lines CHO and HEK-293 or the insect line Sf9 would likely attach and proliferate well based on the findings with the adherent C2C12 line used in the current study.

While the inclusion of leaf material in a food product may prove unpalatable, the relative low density of DCL scaffold compared to the cellular material may be a physical property that enables a simple way to separate the two products. Potentially, following mechanical or chemical disruption in aqueous based solutions, the separated DCL scaffold material would float, and the denser cellular material would sink allowing the cellular material to be easily harvested. Alternatively, others have shown that plant species that are more palatable and commonly consumed are also effective cellular scaffolds (22). We would predict that microbial based decellularization of such species would be achievable based on the findings in this study, a hypothesis that remains to be tested.

## Notes

### Competing Interest Statement

The authors have declared no competing interest.

## References

1. le Coutre J. Editorial: Cultured Meat-Are We Getting it Right? Front Nutr (2021) 8:675797. doi: 10.3389/fnut.2021.675797

2. Broucke K, Pamel V, Coillie V, Herman L, Royen V. Cultured meat and challenges ahead: A review on nutritional, technofunctional and sensorial properties, safety and legislation. Meat Sci (2023) 195:109006. doi: 10.1016/j.meatsci.2022.109006

3. First application for cultivated meat approval in Europe submitted. https://www.foodnavigator.com/Article/2023/07/26/First-application-for-cultivated-meat-approval-in-Europe-submitted [Accessed July 28, 2023]

4. Upside Foods secures USDA approval for its cultivated meat | Reuters. https://www.reuters.com/business/upside-foods-says-receives-label-approval-usda-its-cultivated-meat-2023-06-14/ [Accessed July 28, 2023]

5. Cell-Based Meat One Step Closer to Entering U.S. Market as Two Companies Receive USDA Approvals | Food Safety. https://www.food-safety.com/articles/8690-cell-based-meat-one-step-closer-to-entering-us-market-as-two-companies-receive-usda-approvals [Accessed July 28, 2023]

6. Jones JD, Thyden R, Perreault LR, Varieur BM, Patmanidis AA, Daley L, Gaudette GR, Dominko T. Decellularization: Leveraging a Tissue Engineering Technique for Food Production. (2023) doi: 10.1021/acsbiomaterials.2c01421

7. Bioprocessing by Decellularized Scaffold Biomaterials Q, Zang M, Demitri C, Lu H, Ying K, Shi Y, Liu D, Chen Q. Bioprocessing by Decellularized Scaffold Biomaterials in Cultured Meat: A Review. Bioengineering 2022, Vol 9, Page 787 (2022) 9:787. doi: 10.3390/BIOENGINEERING9120787

8. Allan SJ, Ellis MJ, De Bank PA. Decellularized grass as a sustainable scaffold for skeletal muscle tissue engineering. J Biomed Mater Res A (2021) 109:2471–2482. doi: 10.1002/JBM.A.37241

9. Rybchyn MS, Biazik JM, Charlesworth J, Le Coutre J. Nanocellulose from Nata de Coco as a Bioscaffold for Cell-Based Meat. ACS Omega (2021) 6:33923–33931. doi: 10.1021/ACSOMEGA.1C05235/ASSET/IMAGES/LARGE/AO1C05235_0010.JPEG

10. First hamburger made from lab-grown meat to be served at press conference | Stem cells | The Guardian. https://www.theguardian.com/science/2013/aug/05/first-hamburger-lab-grown-meat-press-conference [Accessed July 26, 2023]

11. Campuzano S, Pelling AE. Scaffolds for 3D Cell Culture and Cellular Agriculture Applications Derived From Non-animal Sources. Front Sustain Food Syst (2019) 3:436742. doi: 10.3389/FSUFS.2019.00038/BIBTEX

12. Ahangar Salehani A, Rabbani M, Biazar E, Heidari Keshel S, Pourjabbar B. The effect of chemical detergents on the decellularization process of olive leaves for tissue engineering applications. Engineering Reports (2023) 5:e12560. doi: 10.1002/ENG2.12560

13. Walawalkar S, Almelkar S. Fabricating a pre-vascularized large-sized metabolically-supportive scaffold using Brassica oleracea leaf. J Biomater Appl (2021) 36:165–178. doi: 10.1177/0885328220968388/ASSET/IMAGES/LARGE/10.1177_0885328220968388-FIG8.JPEG

14. Ahangar Salehani A, Rabbani M, Biazar E, Heidari Keshel S, Pourjabbar B. The effect of chemical detergents on the decellularization process of olive leaves for tissue engineering applications. Engineering Reports (2023) 5:e12560. doi: 10.1002/ENG2.12560

15. Lacombe J, Harris AF, Zenhausern R, Karsunsky S, Zenhausern F. Plant-Based Scaffolds Modify Cellular Response to Drug and Radiation Exposure Compared to Standard Cell Culture Models. Front Bioeng Biotechnol (2020) 8:550667. doi: 10.3389/FBIOE.2020.00932/BIBTEX

16. Wang Y, Dominko T, Weathers PJ. Using decellularized grafted leaves as tissue engineering scaffolds for mammalian cells. In Vitro Cellular and Developmental Biology - Plant (2020) 56:765–774. doi: 10.1007/S11627-020-10077-W/FIGURES/7

17. Emami A, Talaei-Khozani T, Vojdani Z, Zarei fard N. Comparative assessment of the efficiency of various decellularization agents for bone tissue engineering. J Biomed Mater Res B Appl Biomater (2021) 109:19–32. doi: 10.1002/JBM.B.34677

18. Emmanuel E, Keck G, Blanchard JM, Vermande P, Perrodin Y. Toxicological effects of disinfections using sodium hypochlorite on aquatic organisms and its contribution to AOX formation in hospital wastewater. Environ Int (2004) 30:891–900. doi: 10.1016/J.ENVINT.2004.02.004

19. Jho EH, Yun SH, Thapa P, Nam JW. Changes in the aquatic ecotoxicological effects of Triton X-100 after UV photodegradation. Environmental Science and Pollution Research (2021) 28:11224–11232. doi: 10.1007/S11356-020-11362-2/FIGURES/6

20. Olkowska E, Ruman M, Polkowska Z. Occurrence of surface active agents in the environment. J Anal Methods Chem (2014) 2014: doi: 10.1155/2014/769708

21. Ahangar Salehani A, Rabbani M, Biazar E, Heidari Keshel S, Pourjabbar B. The effect of chemical detergents on the decellularization process of olive leaves for tissue engineering applications. Engineering Reports (2023) 5:e12560. doi: 10.1002/ENG2.12560

22. Cheng YW, Shiwarski DJ, Ball RL, Whitehead KA, Feinberg AW. Engineering Aligned Skeletal Muscle Tissue Using Decellularized Plant-Derived Scaffolds. ACS Biomater Sci Eng (2020) 6:3046–3054. doi: 10.1021/ACSBIOMATERIALS.0C00058/ASSET/IMAGES/LARGE/AB0C00058_0005.JPEG

