## Supplementary figures for "Bacterial Decellularization: Non-Chemical Production of Effective Plant Tissue Bio-Scaffolds"

**Rybchyn *et al.*; Supplementary FIGURES**

**FIGURE 1 SUPP**

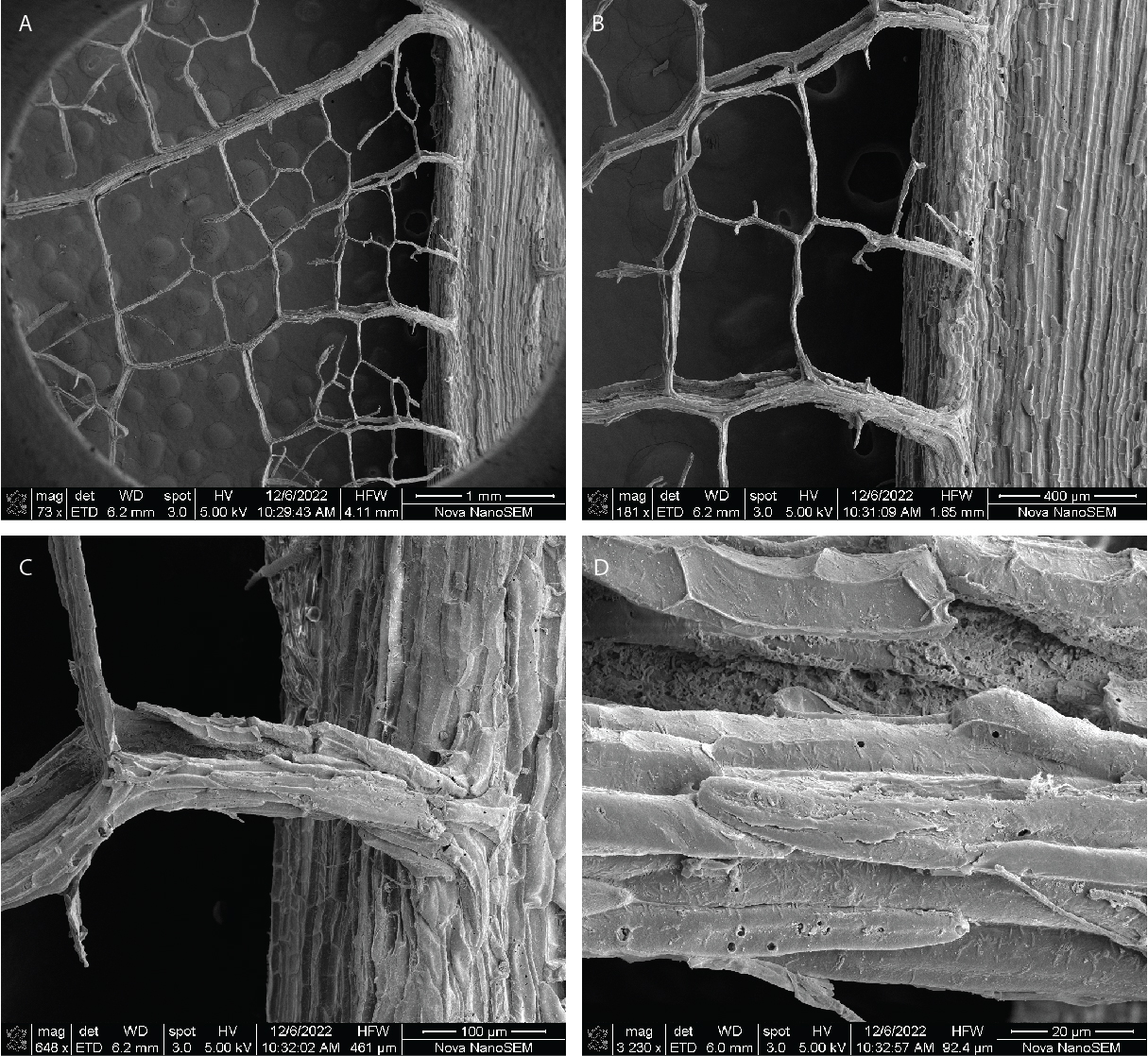

**Figure 1 Supp – Scanning electron microscopy (SEM) revealed that walnut leaves that had been decellularized by bacteria (DCL) comprised structural features that would likely promote myoblast attachment.** (A-D) SEM at increasing magnifications showed that the DCL stem and veins had an uneven surface and were spatially arranged so that individual veins were typically anywhere from 0.1 to 1 mm apart. In some instances, hollow or “pocket” structures were discernable in the DCL, which were presumably where the xylem/phloem vascular bundle originally existed (white arrowhead). Insets are shown (A-C) to show the relative location of increasing magnification. SEM magnifications were as follows (A-73x; B-181x; C-648x; D-3230x).

**FIGURE 2 SUPP**

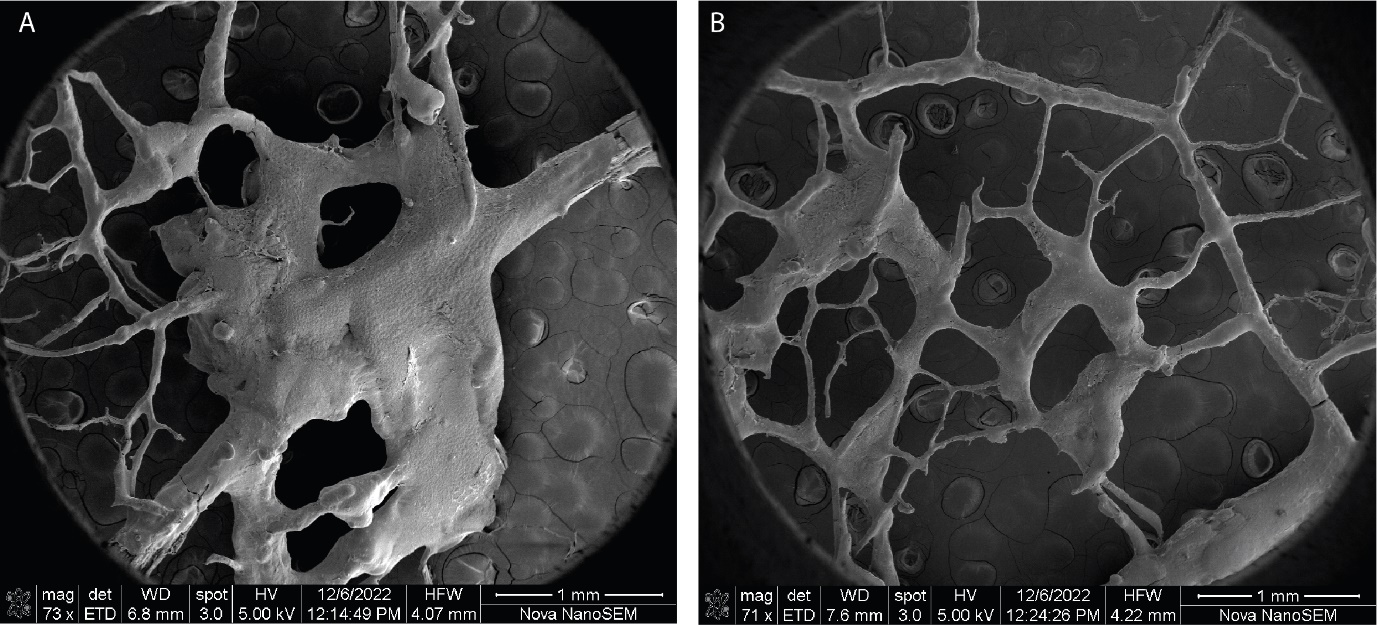

**Figure 2 Supp – C2C12 grew and differentiated extensively over the decellularized leaf (DCL) structure as revealed by low magnification SEM.** (A-B) Low magnification (A-73x; B-71x magnification) SEM images of DCL that was seeded with C2C12 myoblasts and maintained for 3 weeks to promote cellular differentiation. Cellular material is shown as red, DCL without cellular material attached is shown as brown, and the slide background is shown as black.

**FIGURE 3 SUPP**

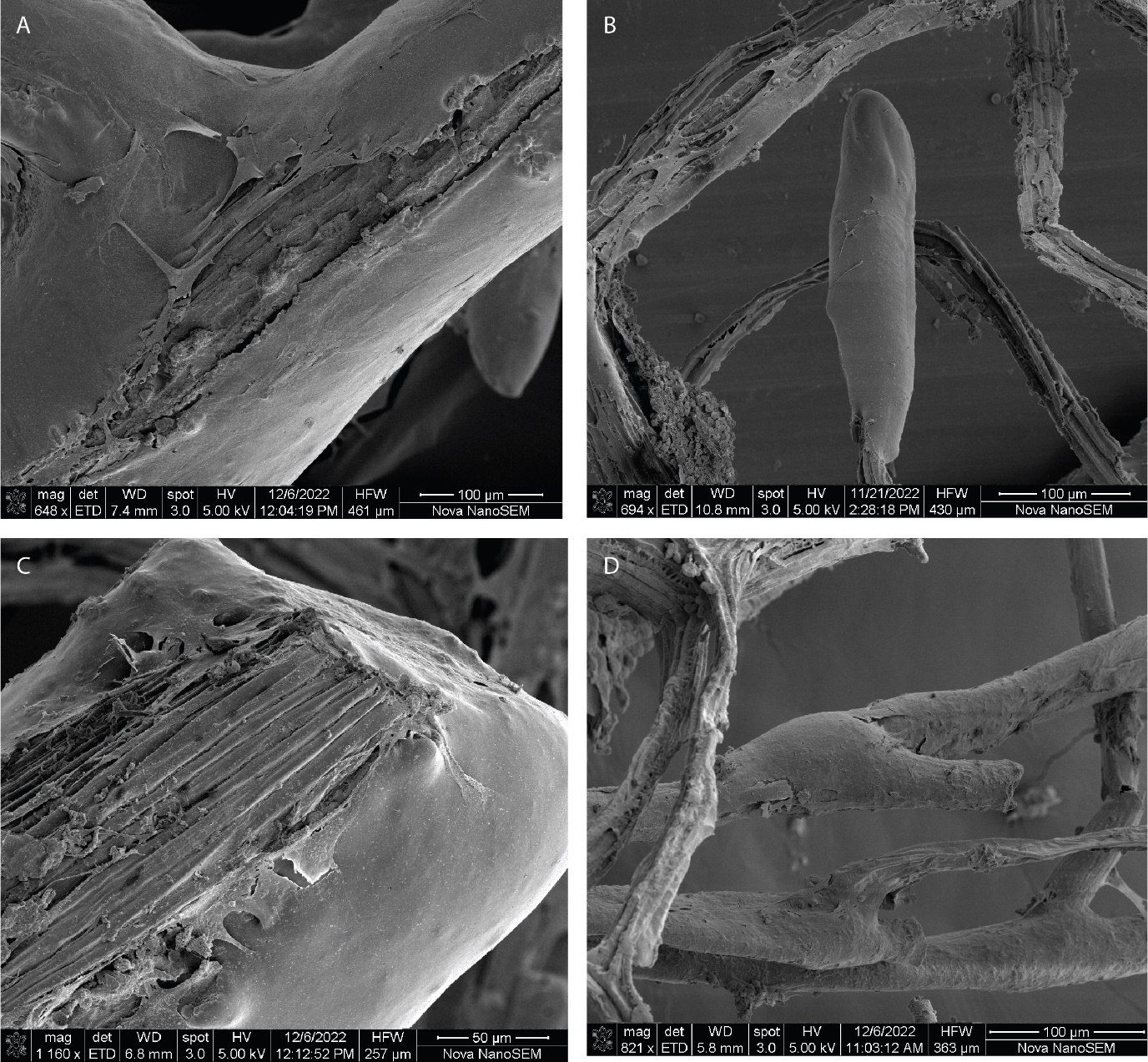

**Figure 3 Supp – Differentiated cellular material grew “tightly” over the veins of the decellularized leaf (DCL) structure as revealed by SEM.** (A-D) SEM images of DCL that was seeded with C2C12 myoblasts and maintained for 3 weeks to promote cellular differentiation. Differentiated cellular material grew tightly over the veins of the DCL. SEM revealed instances where cellular material grew to completely (B and D), or partially (A and C) cover the DCL vein structure.

**FIGURE 4 SUPP**

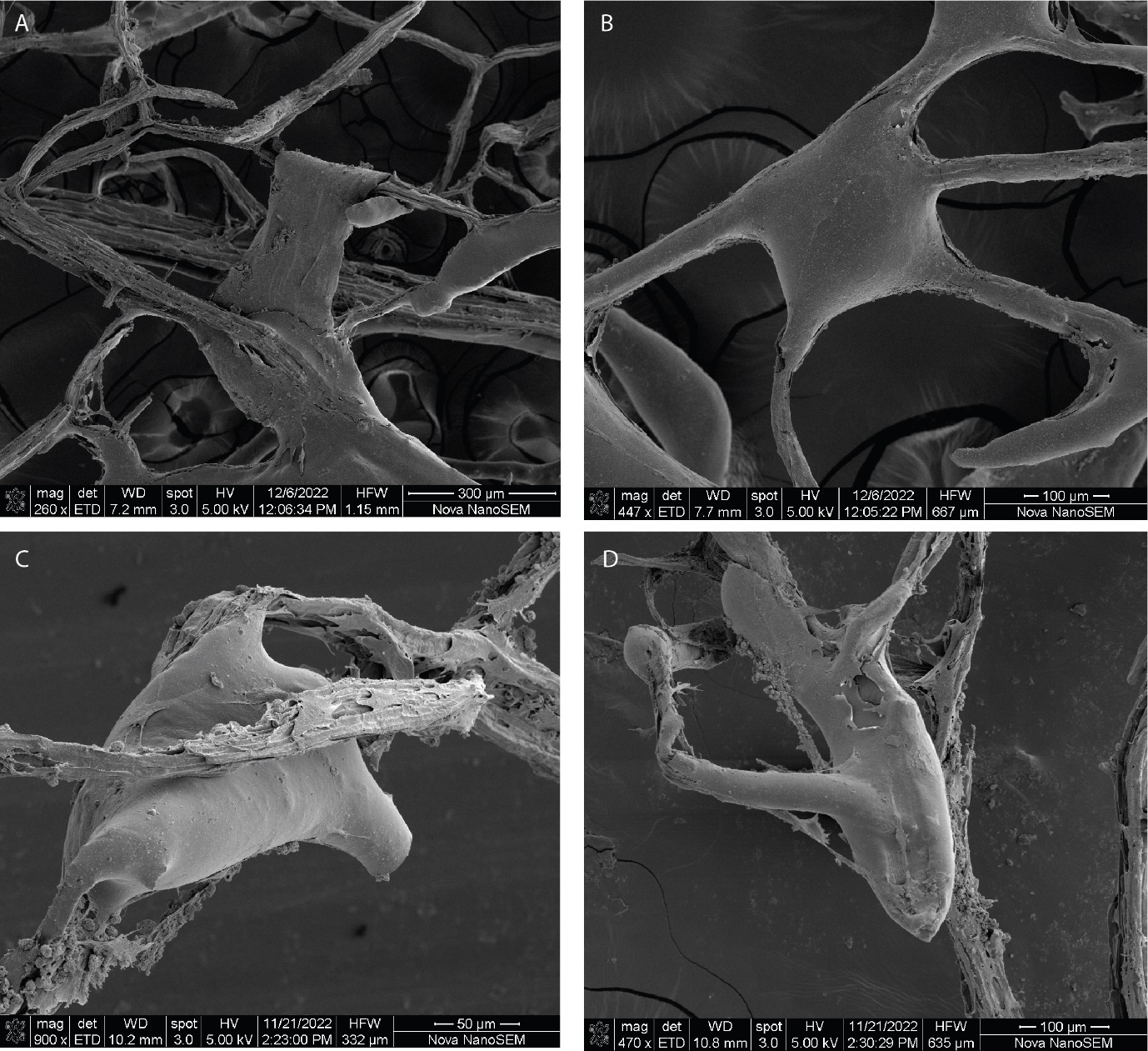

**Figure 4 Supp – Differentiated cellular material grew between the individual veins of the decellularized leaf (DCL) structure as revealed by SEM.** (A-D) SEM images of DCL that was seeded with C2C12 myoblasts and maintained for 3 weeks to promote cellular differentiation. Differentiated cellular material grew between the veins of the DCL producing novel cellular structures. SEM revealed instances where; in (A) a novel differentiated cellular structure grew to be supported at either end by the DCL, as shown by the “looping” of the cellular material over the DCL (A, white arrow); in (B) a completely novel differentiated cell structure was detected growing between the DCL veins with a central mass (B, white arrow) and “arms” extending to the DCL veins to support the structure; in (C) where the mass of a cellular growth attached to one DCL vein (C, white arrow), was sterically supported by a second vein (C, green arrow); in (D) where a cellular growth was supported by several attachment points to DCL veins and grew in to a large supported mass (D, white arrow).

**FIGURE 5 SUPP**

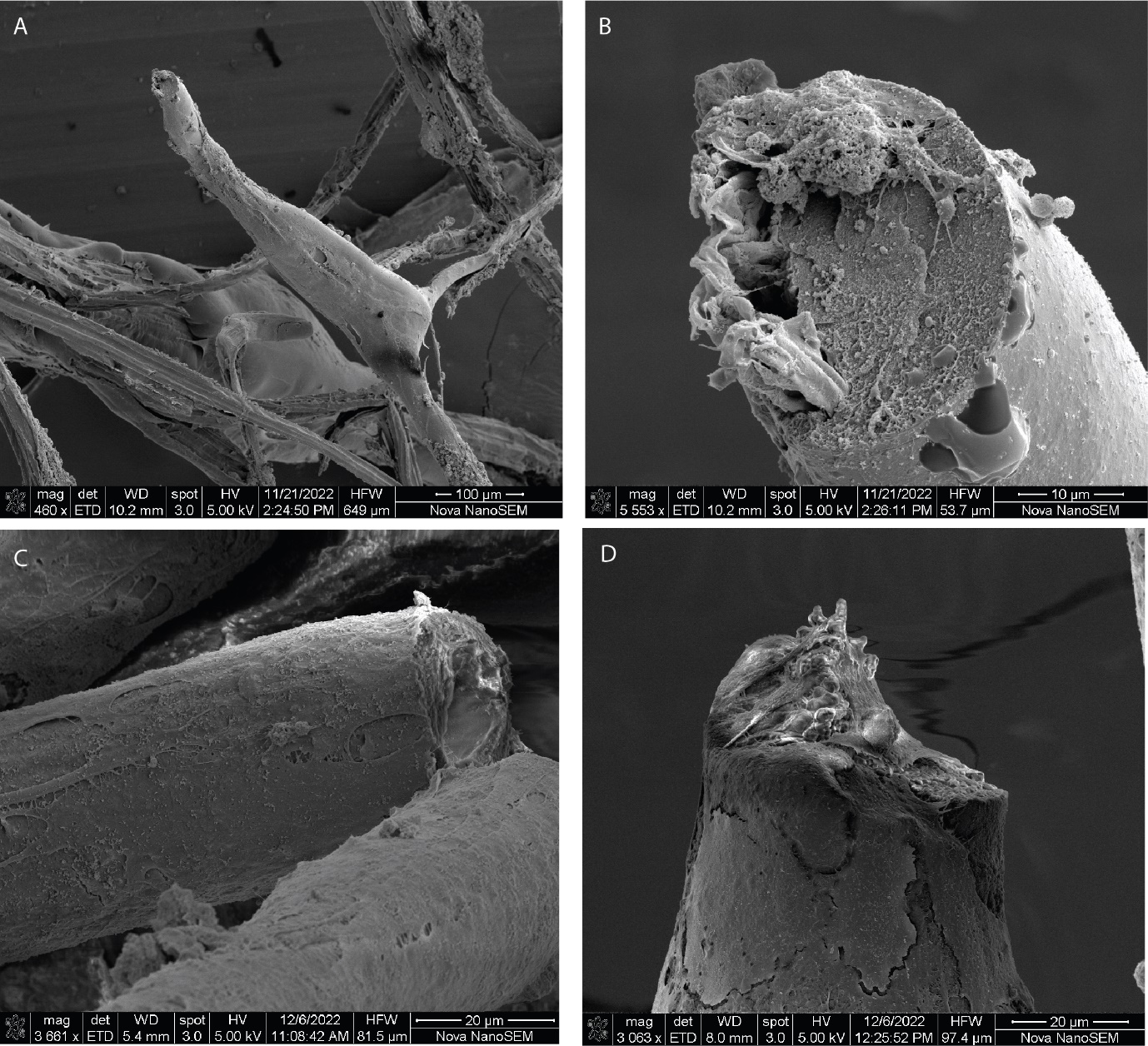

**Figure 5 Supp - Scanning electron microscopy (SEM) revealed novel differentiated fibril structures were growing from anchor points on the DCL.** (A-D) SEM images of DCL that was seeded with C2C12 myoblasts and maintained for 3 weeks to promote cellular differentiation. SEM revealed novel myofibril structures that extended from anchor points on the DCL scaffold. (A) Novel cylindrical myofibril structures (white arrow) extended from anchor points on the DCL (green arrow). (B) A high SEM magnification inset of (A) and (D, a different fibril structure) show active cellular remodeling at the end of the fibril, revealing that myofibrils were still actively forming at the point of fixation (3 weeks post-seeding). The interior of the ends of these actively forming fibrils was solid. (C) SEM of a fractured pre-formed fibril structure revealed a hollow interior structure (white arrow) and thick cellular outer layers (green arrow) which differed from the active point of fibril formation shown in B and D.
